# Viability markers for determination of desiccation tolerance and critical stages during dehydration in *Selaginella* species

**DOI:** 10.1101/2021.03.23.436672

**Authors:** Gerardo Alejo-Jacuinde, Tania Kean-Galeno, Norma Martínez-Gallardo, J. Daniel Tejero-Díez, Klaus Mehltreter, John P. Délano-Frier, Melvin J. Oliver, June Simpson, Luis Herrera-Estrella

**Affiliations:** National Laboratory of Genomics for Biodiversity (Langebio), Centro de Investigación y de Estudios Avanzados del Instituto Politécnico Nacional, 36824, Irapuato, Guanajuato, Mexico; Department of Genetic Engineering, Centro de Investigación y de Estudios Avanzados del Instituto Politécnico Nacional, 36824 Irapuato, Guanajuato, Mexico; Institute of Genomics for Crop Abiotic Stress Tolerance (IGCAST), Department of Plant and Soil Science, Texas Tech University, 79409 Lubbock, Texas, USA; Department of Biotechnology and Biochemistry, Centro de Investigación y de Estudios Avanzados del Instituto Politécnico Nacional, 36824 Irapuato, Guanajuato, Mexico; Facultad de Estudios Superiores Iztacala, Universidad Nacional Autónoma de México, 54090 Tlalnepantla, Estado de Mexico, Mexico; Red de Ecología Funcional, Instituto de Ecología A.C., 91070 Xalapa, Veracruz, México; Division of Plant Sciences, Interdisciplinary Plant Group, University of Missouri, Columbia, MO 65211, USA

**Keywords:** critical stage, desiccation tolerance, Selaginella, tissue viability.

## Abstract

Plants can tolerate some degree of dehydration but below a threshold of water content most plants die. However, some species display specific physiological, biochemical, and molecular responses that allow survival to desiccation. Some of these responses are activated at critical stages during water loss and could represent the difference between desiccation tolerance (DT) and death. Here, we report the development of a simple and reproducible system to determine DT in *Selaginella* species. This system is based on the use of excised tissue (explants), exposed to a dehydration agent inside small containers, rather than whole plants making it faster, better controlled, and potential use under field conditions. We also report that the triphenyltetrazolium chloride test is a simple and accurate assay to determine tissue viability. The explant system is particularly useful to identify critical points during the dehydration process and was applied to identify novel desiccation-tolerant *Selaginella* species. Our data suggest that desiccation-sensitive *Selaginella* species have a change in viability when dehydrated to 40% RWC, indicating the onset of a critical condition at this water content. Comparative studies at these critical stages could provide a better understanding of DT mechanisms and unravel insights into the key responses to survive desiccation.

**Highlight:** In this article, we developed a simple and efficient system to determine desiccation tolerance and critical stages during the dehydration process in *Selaginella* that can be applied to other plant species.

## Introduction

Desiccation tolerance (DT) is defined as the ability of an organism to dry to equilibrium with moderately dry air (50% or lower relative humidity) and to resume its metabolism when rehydrated (Alpert and Oliver, 2002; Alpert, 2005). Drying cells to 50% relative humidity (RH), corresponding to a water potential of about -100 MPa, leads to metabolic arrest since cellular water content is below the level required to form an aqueous monolayer around macromolecules (Alpert, 2005; Leprince and Buitink, 2015). Essential features of DT include the ability of plants or tissues to limit the damage to a repairable level, to maintain cellular and subcellular physiological integrity in the dry state, and to repair damage upon rehydration (Bewley, 1979; Alpert and Oliver, 2002). In most basal clades of land plants (bryophytes), DT exhibits a high degree of plasticity and is dependent upon external and internal environmental factors (Oliver *et al*., 2020). Nevertheless, many bryophytes exhibit DT that is characterized by constitutive protection mechanisms that allow them to survive desiccation regardless of the drying rate, whereas in tracheophytes DT is mainly based on the activation of protection mechanisms that lead to DT only if water loss is gradual (Oliver *et al*., 2000).

During dehydration plant cell volume can decrease by 60 to 80%, therefore, to survive the mechanical stress caused by water loss, cells of desiccation-tolerant pteridophytes and eudicot angiosperms accumulate compatible solutes, increase vacuolation and activate cell wall folding (Farrant *et al*., 2007; Oliver *et al*., 2020). Sensitive species do not possess these mechanisms and sub-cellular damage, for example plasma membrane disruption, is lethal (Farrant *et al*., 2007). Accumulation of soluble sugars is directly related to DT in seeds and vegetative tissues (Alpert and Oliver, 2002), and their proposed functions include water replacement in membranes and macromolecules, filling and stabilization of vacuoles, and vitrification (a glassy state of the cytoplasm) (Hoekstra *et al*., 2001; Dinakar and Bartels, 2013). The principal protective sugar that accumulates to high amounts during drying is sucrose, however carbohydrate metabolism is highly diverse among desiccation-tolerant plants and other molecules such as trehalose, raffinose family oligosaccharides, and even unusual sugars such as octulose can also accumulate in tolerant species (Zhang *et al*., 2016).

Although desiccation-tolerant species shut down photosynthetic activity during early dehydration, electron flow in light-harvesting reactions continues resulting in overproduction of reactive oxygen species (ROS), which can lead to damage to macromolecules (Dinakar *et al*., 2012; Challabathula *et al*., 2018). Mechanisms to limit ROS production in desiccation-tolerant species include morphological changes such as leaf folding to minimize the area exposed to light, preferential exposure of the leaf abaxial surface, stem curling, and shading (Lebkuecher and Eickmeier, 1991; Bartels and Hussain, 2011). Desiccation-tolerant species are classified as homoiochlorophyllous or poikilochlorophyllous, according to the strategy to either protect or dismantle their photosynthetic machinery during dehydration (Challabathula *et al*., 2018). Poikilochlorophyllous species evolved the ability to dismantle thylakoid membranes and degrade chlorophyll in order to reduce ROS production in the desiccated state; these components are then resynthesized and reassembled during rehydration (Dinakar *et al*., 2012). In contrast, homoiochlorophyllous plants retain and protect their photosynthetic apparatus during desiccation for a rapid reactivation upon rehydration (Dinakar *et al*., 2012). Thus, homoiochlorophyllous species require more effective mechanisms of protection against ROS in comparison to poikilochlorophyllous species (Farrant *et al*., 2007; Challabathula *et al*., 2016; Georgieva *et al*., 2017).

Desiccation associated damage is lethal for most plants and the understanding of how protective mechanisms in desiccation-tolerant species allow them to survive extreme dehydration will be important for the future breeding of drought-tolerant crops. Furthermore, to gain a better understanding of DT it is necessary to determine accurate indicators to define viability and critical stages during the desiccation process. The establishment of these indicators was carried out in *Selaginella* species, a cosmopolitan group which occupies a broad diversity of habitats ranging from tropical rainforests to deserts (Zhou *et al*., 2015). Here we report an explant dehydration system in conjunction with methodologies to measure tissue damage and recovery after a desiccation process. The implemented methods allowed to determine the type of DT response that can showed desiccation-tolerant *Selaginella* species and the specific water contents at which a sensitive species loses tissue viability. Our methodology represents a simple and robust tool to determine DT ability and critical stages during the dehydration process.

## Materials and methods

### Plant material

A total of 16 *Selaginella* species were included in the present study, a list of species and collection sites is provided in **Supplementary Table S1**. Specimens were taxonomically determined according to the key in Mickel & Smith (2004) and voucher specimens were deposited at MEXU herbarium, UNAM. Plants were transferred to pots or trays (depending on the plant size). Tolerant species *S. lepidophylla* and *S. sellowii* were maintained in the dried state at room temperature until used. Sensitive species *S. denticulata* and *S. silvestris* were maintained in a growth chamber (22 °C, 16 h light period). All *Selaginella* samples used to determine DT ability were maintained in hydrated conditions in a greenhouse under ambient light conditions until use.

### Explant desiccation treatments

Desiccation-tolerant species (*S. lepidophylla* and *S. sellowii*) were watered and maintained in a hydrated condition in a growth chamber (22 °C, 16 h light period) for at least 5 days before DT experiments. Desiccation treatments involving explants placed in small closed containers are described in detail in the **Supplementary Protocol S1**. The desiccated state was evaluated 1 week after explants were placed in the drying system (25 °C, 16 h light period). The rehydration state was evaluated by adding 5 ml of deionized water to desiccated tissue and analyzing it after 24 h.

### Electrolyte leakage measurements

Explants with an initial fresh weight of 100 ± 1 mg were briefly rinsed in deionized water to remove the contents of cut cells and excess water was removed by blotting. Desiccated explants were transferred to 50 ml polypropylene tubes and rehydrated. When the rehydration step was completed, the tubes were topped up to exactly 25 ml with deionized water and incubated for 4 h at room temperature in an orbital shaker (160 rpm). Well-watered samples were included to evaluate explants under hydrated conditions. Electrolyte leakage (C_i_) was measured with a HI2003 conductivity meter (Hanna Instruments, USA). Subsequently, samples were placed in boiling water for 30 min. After cooling, the conductivity was measured and this value represents the total amount of ions in the sample (C_max_). The conductivity of deionized water was also measured (C_w_) and Relative Electrolyte Leakage (REL) was calculated using the following formula:

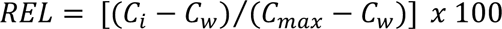

### Chlorophyll content and maximum quantum efficiency

Explants for chlorophyll quantification were transferred to 15 ml polypropylene tubes, frozen in liquid nitrogen, and stored at -80 °C until use. Chlorophyll extraction and quantification (total chlorophyll, chlorophyll *a*, and chlorophyll *b*) were carried out according to Richardson *et al*. (2002). The incubation time with DMSO (65 °C) was modified to 45 min for the tolerant species. Chlorophyll quantification was adapted to measure in 96-well plates.

Maximum quantum efficiency of PSII (Fv/Fm) was determined in 15 min dark-adapted explants with a Pocket PEA chlorophyll fluorimeter (Hansatech Instruments, UK).

### Antioxidant potential assays, total phenol and flavonoid content

Explants with an initial fresh weight of 100 ± 1 mg were used for the following protocols. Tissue was transferred to 2 ml polypropylene tubes containing a steel bead, frozen in liquid nitrogen, and stored at -80 °C until use. Frozen tissue was ground using a TissueLyser II (Qiagen Inc., USA) with 3 rounds of shaking at 30 Hz for 30 s. *In vitro* antioxidant activity was measured by the ferric-ion reducing antioxidant power (FRAP) and the 2,2-diphenyl-1-picrylhydrazyl (DPPH) assays according to the methodology described by Yahia *et al*. (2011). Total phenol compound (TPC) content was determined according to Maranz *et al*. (2003). Total flavonoid (TF) content was determined according to Sakanaka *et al*. (2005). All procedures were adapted to measure in 96-well plates.

### RNA integrity

Tissue was transferred to 2 ml polypropylene tubes containing a steel bead, frozen in liquid nitrogen, and stored at -80 °C until use. Frozen tissue was ground using a TissueLyser II (Qiagen Inc., USA) with 3 rounds of shaking at 30 Hz for 30 s. RNA was extracted from frozen ground tissue using PureLink™ Plant RNA Reagent (Invitrogen) according to the manufacturer’s instructions. RNA integrity was examined by 1.2% agarose gel electrophoresis.

Determination of tissue viability using the triphenyltetrazolium chloride (TTC) staining was adapted from Lin *et al*. (2001) and Ruf & Brunner (2003). The methodology to determine viability by TTC assay and image analysis to measure the proportion of the viable area is described in detail in **Supplementary Protocol S1**. Image processing was carried out using GIMP software (version 2.10.12) and pixel quantification to determine viable explant areas using the software Easy Leaf Area (Easlon and Bloom, 2014).

### Whole plant desiccation treatments

Greenhouse whole plant experiments were carried out by withholding water from plants for 30 days, including at least 3 individuals (pots or trays) per species. Plants were watered and evaluated 2 days after rehydration. Species in which at least half of the tissue of each individual retained their green color at rehydration were classified as desiccation-tolerant.

### Statistical analysis

Results are expressed as mean values ± standard deviation; the number of replicates is given in figure legends. The significance of differences between control (hydrated, unstressed tissue) and each treatment was analyzed using Tukey’s HSD test.

## Results

### Drying rate establishment and evaluation by physical appearance

Desiccation experiments were carried out in a simple system based on explants, rather than whole plants, to evaluate DT-related traits in different *Selaginella* species (**Fig. 1A**). Explant desiccation treatments (and technical replicates) were performed in individual desiccators for each condition in order to obtain reproducible results. The drying system comprises a small plastic container, the bottom part of a Petri dish to place the explants inside the container and a dehydration agent (saturated salt solutions or silica gel; **Fig.1B**) as described in detail in **Supplementary Protocol S1**. Explants of desiccation-tolerant *Selaginella* species did not survive if drying is too fast to allow the acquisition of DT (e.g., drying in an oven at 65 °C). To determine appropriate rates of water loss, we evaluated three drying regimes: slow drying using a saturated solution of KNO_2_ (45.5-46.5% RH), moderate drying using a saturated solution of MgCl_2_ (32-33% RH), and rapid drying using silica gel (5-7% RH). To test the drying regimes, we used explants from two well characterized desiccation-tolerant species *S. lepidophylla* and *S. sellowii.* The explants of these two tolerant species recovered their initial physical appearance upon rehydration after the desiccation process (green color; slight oxidation in some explants during fast drying using silica gel) using the three types of dehydration agents. Relative electrolyte leakage (REL) was lower for all drying rates in *S. sellowii* in comparison to *S. lepidophylla* (**Fig. 1C**), indicating better membrane protection mechanisms in the former of the two desiccation-tolerant species. Measurement of REL showed that the moderate drying rate produces the lowest membrane damage in both tolerant species (**Fig. 1C**). Therefore, the moderate drying rate (MgCl_2_) which equilibrates at around -149 MPa (Xiao *et al*., 2018), a stricter water potential than the threshold for determination of plant DT (-100 MPa) was selected for the comparisons between desiccation-tolerant and sensitive species. We tested whether the explant system could be used to differentiate between desiccation-tolerant and sensitive species. With this aim, we subjected explants from the same two desiccation-tolerant species (*S. lepidophylla* and *S. sellowii*) and two previously characterized sensitive species (*S. silvestris* and *S. denticulata*) to desiccation. Both tolerant and sensitive *Selaginella* species reached equilibrium (desiccated state) within 48 h after explants were placed in the drying system (**Supplementary Fig. S1**). Evaluation of physical appearance of explants showed complete recovery after rehydration of tolerant species *S. lepidophylla* and *S. sellowii*, whereas the explants of the sensitive species suffered severe oxidation (**Fig. 2A**). Tissue of desiccation-tolerant species showed ordered morphological changes during desiccation, including compact stem curling and/or microphyll folding, whereas microphylls of sensitive species underwent shrinkage but most of the green tissue (photosynthetic area) in desiccated state remained exposed to light (**Fig. 2A**).

**Figure 1.**
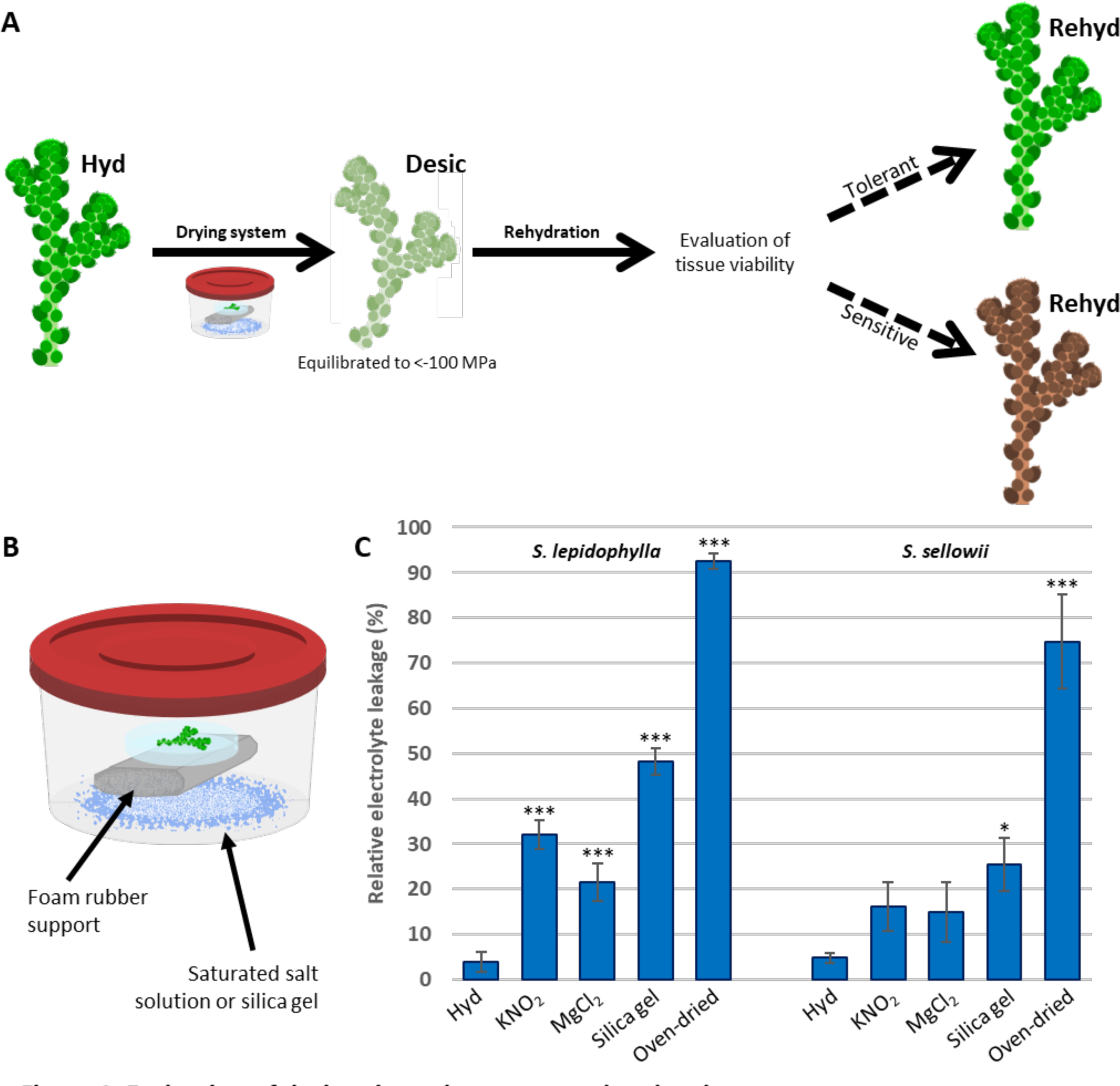
Evaluation of desiccation tolerance at explant level. (A) Schematic diagram of experimental design using explants. Evaluated stages included hydrated (Hyd), desiccated (Desic) and rehydrated (Rehyd) conditions. (B) Representation of the drying system; the positions of the tissue (open Petri dish over a support) and a humidity control agent are depicted. (C) Comparison of relative electrolyte leakage (REL) of *S. lepidophylla* and *S. sellowii* explants exposed to different drying rates produced by saturated salt solutions or silica gel. Hydrated explants (Hyd) were used as control of unstressed tissue. Oven-dried explants (65 °C) represent a state of irreparable damage. Each bar represents the mean of 3 replicates and error bars indicate standard deviation. Bars with asterisks are significantly different from Hyd (Tukey’s HSD test, \**P*>0.05, \*\**P*>0.01, \*\*\**P*>0.001).

**Figure 2.**
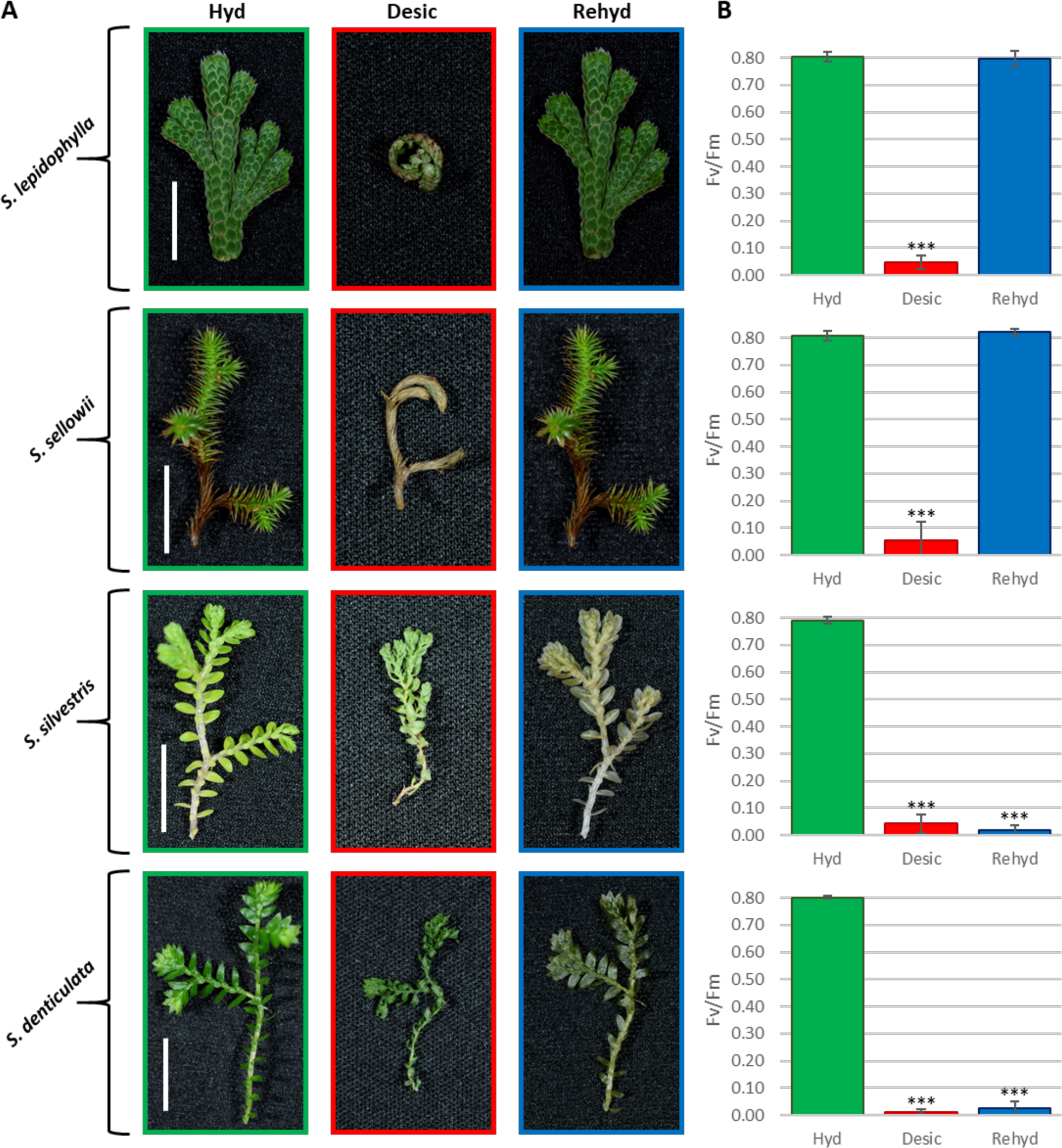
Evaluation of explant recovery by visual inspection and determination of quantum efficiency of PSII. (A) Physical appearance of *S. lepidophylla*, *S. sellowii*, *S. silvestris* and *S. denticulata* explants during a dehydration-rehydration cycle. (B) Maximum quantum efficiency of PSII (Fv/Fm). Hydration stage is color-coded: green, red and blue indicate hydrated (Hyd), desiccated (Desic) and rehydrated (Rehyd), respectively. Error bars represent a standard deviation of 10 points per condition. Bars with asterisks are significantly different from Hyd (Tukey’s HSD test, \*\*\**P*>0.001). Scale bar = 1 cm

### Viability evaluation through homoiochlorophyllous properties

Retention of the photosynthetic apparatus in tolerant *Selaginella* species allowed the implementation of maximum quantum efficiency of PSII (Fv/Fm) measurements to evaluate recovery after desiccation (**Fig. 2B**). Both tolerant and sensitive *Selaginella* species showed a significant decrease in Fv/Fm values after desiccation, but only tolerant plants recovered the initial values upon rehydration (similar to hydrated, unstressed controls) (**Fig. 2B**). Complete recovery of Fv/Fm values could be associated with an effective protection of the photosynthetic apparatus, including pigments. Chlorophyll fluorescence, evaluated by photographs of tissue exposed to UV light, showed that for desiccation-tolerant species fluorescence was similar before and after desiccation, whereas for sensitive plants a diminished fluorescence was observed after desiccation (**Supplementary Fig. S2**). Quantification of total chlorophyll contents determined that *S. lepidophylla* and *S. sellowii* showed a decrease to between 82 and 80.5% of the hydrated total chlorophyll content respectively in the desiccated state, but recovered to 92.3 and 94.3% of the hydrated total chlorophyll content respectively upon rehydration (**Fig. 3A**). In contrast to the desiccation-tolerant plants, the sensitive species *S. silvestris* and *S. denticulata* showed a decrease to 68.6 and 79.6% of the hydrated total chlorophyll content respectively at desiccation, and a further reduction to 52.8 and 46.1% of the hydrated total chlorophyll content respectively after rehydration (**Fig. 3A**).

**Figure 3.**
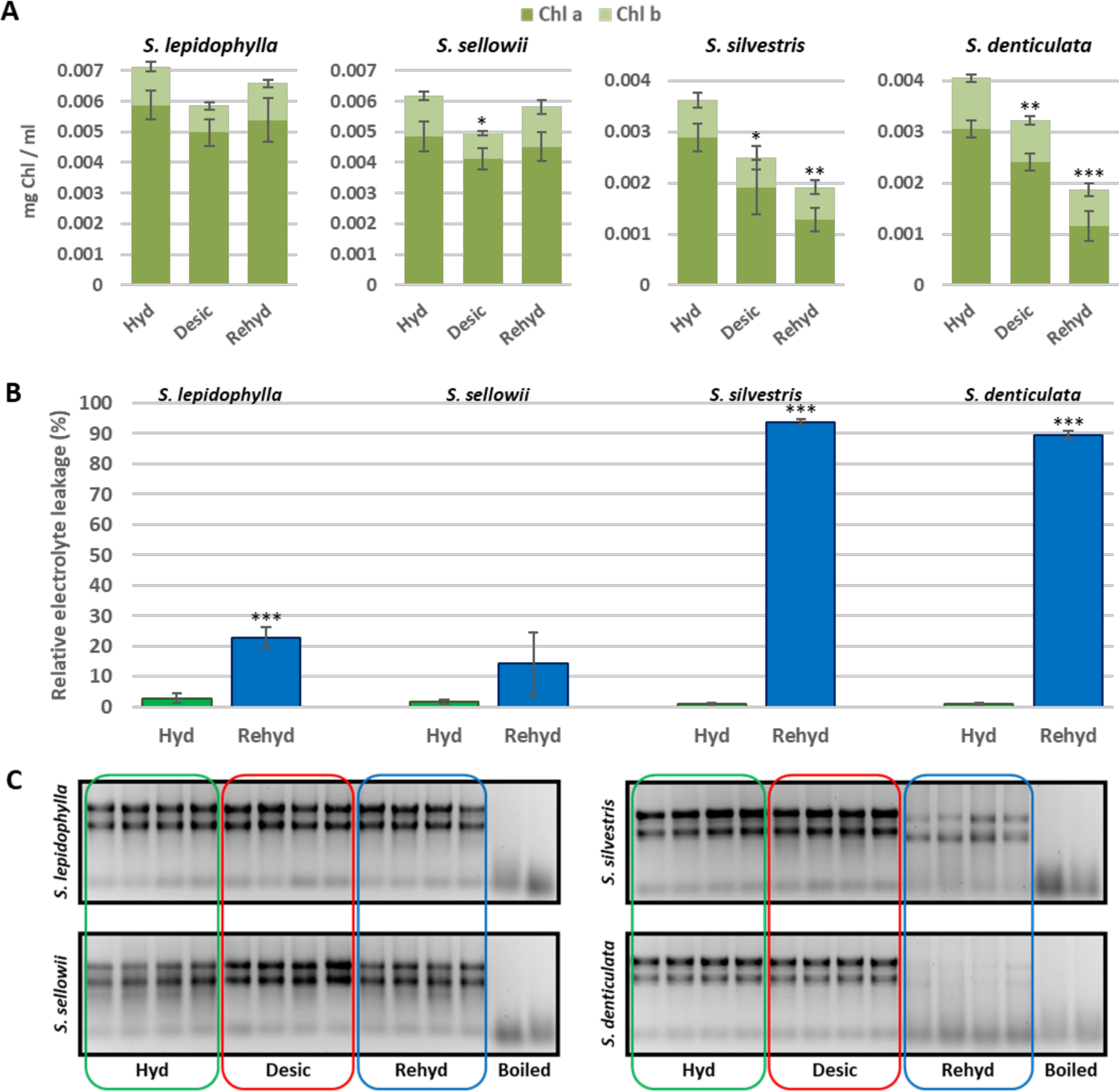
Maintenance of chlorophyll content and integrity of essential components during the desiccation process. (A) Quantification of chlorophyll *a* (dark green) and chlorophyll *b* (light green) under hydrated (Hyd), desiccated (Desic), and rehydrated (Rehyd) conditions. (B) Membrane damage indicated by relative electrolyte leakage before and after desiccation. (C) Ribosomal RNA integrity examined by agarose gel electrophoresis. Boiled explants were included as non-viable tissues. Error bars represent standard deviation of 4 replicates. Bars with asterisks are significantly different from Hyd (Tukey’s HSD test, \**P*>0.05, \*\**P*>0.01, \*\*\**P*>0.001).

Antioxidant potential was evaluated by ferric-ion reducing antioxidant power (FRAP) and 2,2-diphenyl-1-picrylhydrazyl (DPPH) assays (**Table 1**). In hydrated conditions, tolerant species exhibit at least a two-to four-fold higher level of antioxidant capacity in both assays. Although both tolerant and sensitive species increased their antioxidant capacity in response to desiccation, the levels observed in the tolerant species *S. lepidophylla* and *S. sellowii* remained significantly higher than in the sensitive plants (**Table 1**). During rehydration, antioxidant potential in sensitive species decreased to even lower levels than hydrated tissue, whereas in the tolerant species antioxidant potential remained higher during rehydration than in the initial hydrated state (**Table 1**), but only the FRAP assay showed this behavior for explants of *S. sellowii*. Additionally, total phenol content (TPC) and total flavonoids (TF) were determined and displayed similar patterns seen in the antioxidant potential assays (**Table 1**).

**Table 1.** Differences in antioxidant potential between tolerant and sensitive species.

| Species | Condition | FRAP<br>(mg AA/g FW) | DPPH<br>(mg AA/g FW) | TPC<br>(mg/g FW) | TF<br>(mg/g FW) |
| --- | --- | --- | --- | --- | --- |
| *S. lepidophylla* | Hydrated | 1.21 (0.49) | 0.63 (0.37) | 6.93 (1.67) | 3.60 (1.16) |
| *S. lepidophylla* | Desiccated | 5.82 (0.77) | 6.70 (0.42) | 12.90 (2.37) | 9.30 (3.13) |
| *S. lepidophylla* | Rehydrated | 2.23 (0.35) | 1.18 (0.28) | 8.44 (0.40) | 4.71 (1.66) |
| *S. sellowii* | Hydrated | 3.43 (0.31) | 4.40 (0.23) | 14.98 (1.46) | 7.42 (1.70) |
| *S. sellowii* | Desiccated | 5.06 (0.60) | 7.91 (0.90) | 18.61 (2.18) | 10.04 (2.42) |
| *S. sellowii* | Rehydrated | 4.04 (0.96) | 3.07 (1.15) | 12.74 (2.39) | 6.57 (2.89) |
| *S. silvestris* | Hydrated | 0.61 (0.16) | 0.07 (0.02) | 3.34 (0.37) | 1.62 (0.19) |
| *S. silvestris* | Desiccated | 0.93 (0.18) | 0.26 (0.07) | 4.32 (0.50) | 2.20 (0.35) |
| *S. silvestris* | Rehydrated | 0.23 (0.09) | -0.05 (0.01) | 2.22 (0.21) | 1.28 (0.11) |
| *S. denticulata* | Hydrated | 0.74 (0.12) | -0.02 (0.01) | 1.60 (0.12) | 1.80 (0.32) |
| *S. denticulata* | Desiccated | 1.78 (0.19) | 0.90 (0.06) | 3.70 (0.32) | 3.39 (0.58) |
| *S. denticulata* | Rehydrated | 0.61 (0.18) | -0.11 (0.02) | 1.44 (0.17) | 1.56 (0.20) |

Antioxidant potential measured by FRAP (Ferric-ion Reducing Antioxidant Power) and DPPH (2,2-diphenyl-1-picrylhydrazyl) assays expressed as ascorbic acid (AA) equivalents. TPC (Total Phenol Content) and TF (Total Flavonoids) contents are expressed as caffeic acid and catechin equivalents respectively. Data represent the mean value of 4 replicates, data in brackets indicates standard deviation. Fresh Weight (FW).

### Retention and integrity of essential components

Tissue of tolerant *Selaginella* species apparently did not exhibit damage at the rehydration stage when evaluated by visual inspection (**Fig. 2A**). However, measurements of REL indicated some degree of membrane damage in tolerant plants (**Fig. 3B**). On the other hand, membrane integrity in the sensitive species (*S. silvestris* and *S. denticulata*) is completely compromised upon rehydration (around 90% of REL; **Fig. 3B**).

Presence of ribosomes is required to reinitiate protein biosynthesis at rehydration, therefore, we tested whether ribosomal RNA (rRNA) integrity could also be informative about the survival of plants under desiccation. Integrity of rRNA was maintained in all *Selaginella* species at desiccated state, at least during the short period evaluated (1 week). However, rRNA was clearly degraded in the sensitive species during rehydration, whereas integrity of rRNA was largely intact in tolerant species (**Fig. 3C**).

### Determination of tissue viability by analysis of respiratory chain activity

Triphenyltetrazolium chloride (TTC) is commonly used to assess seed viability (Lopez Del Egido *et al*., 2017). The activity of the mitochondrial respiratory chain reduces TTC to an insoluble red compound (triphenylformazan), indicating that the tissue is metabolically active. Therefore, we tested whether TTC could be used to determine tissue viability after a desiccation process. Fresh tissue showed a strong red color indicating that the implemented TTC test can be used to evaluated viability in vegetative tissues. Explants were then tested using the TTC assay following a desiccation treatment (**Fig. 4**). Rehydrated explants of desiccation-tolerant species showed a strong red color similar to unstressed tissue. In contrast, rehydrated explants of sensitive species had no detectable TTC reduction (similar to that observed in control explants killed by boiling), indicating that these explants were not metabolically active (do not recover respiration) after the dehydration/rehydration cycle (**Fig. 4**).

**Figure 4.**
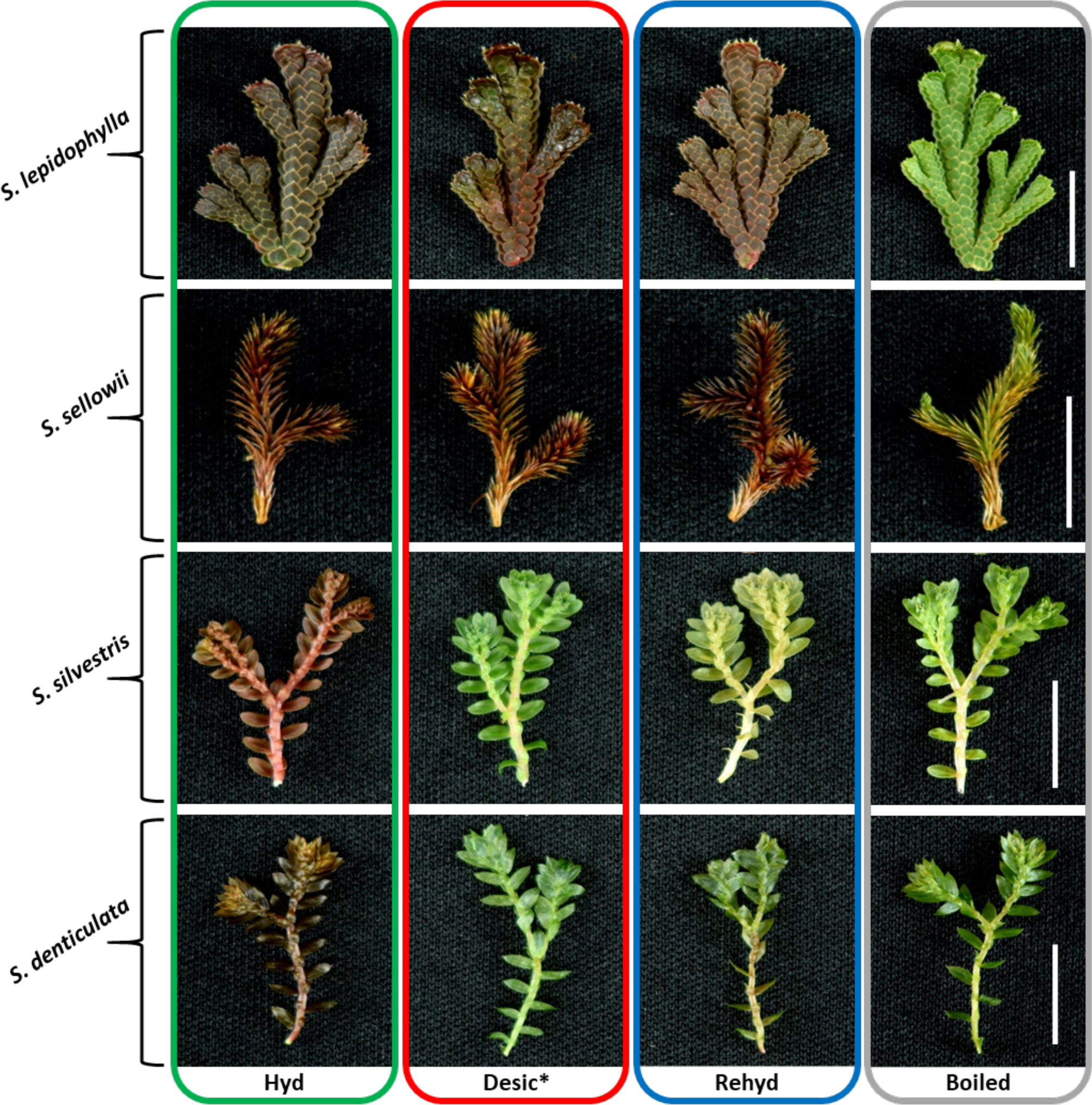
Determination of tissue viability based on respiratory activity. The respiratory chain produces an insoluble pink/red compound by the reduction of triphenyltetrazolium chloride (TTC). Desiccated tissue was directly submerged in TTC solution (Desic*). No TTC reduction was observed in boiled explants included as non-viable tissues (gray box). Abbreviations and color code is similar to Figure 2. Scale bar = 1 cm

### Identification of novel desiccation-tolerant Selaginella species

Determination of tissue viability by TTC test resulted in an efficient and accurate methodology that was applied to classify previously uncharacterized *Selaginella* species as desiccation-tolerant or sensitive (**Fig. 5**). Based on the TTC staining results, the following species were classified as desiccation-tolerant: *S. extensa* (**Fig. 5A**), *S. rupincola* (**Fig. 5B**), *S. wrightii* (**Fig. 5C**), *S. nothohybrida* (**Fig. 5D**), *S. ribae* (**Fig. 5F**), *S. polyptera* (**Fig. 5G**), and *S. schiedeana* (**Fig. 5H**). The analysis also included a sample of *S. pilifera* (**Fig. 5E**), a previously reported desiccation-tolerant species (Eickmeier, 1980). Explants of the samples *S. flexuosa* (**Fig. 5J**), *S. lineolata* (**Fig. 5K**), and *S. stellata* (**Fig. 5L**) were classified as desiccation-sensitive species by this analysis. A different staining pattern was observed in *S. delicatissima* (**Fig. 5I**) in which only the apices of some explants (in one-third of the evaluated explants) remained viable after the desiccation process, therefore, this species was classified as a tissue-specific tolerant species.

**Figure 5.**
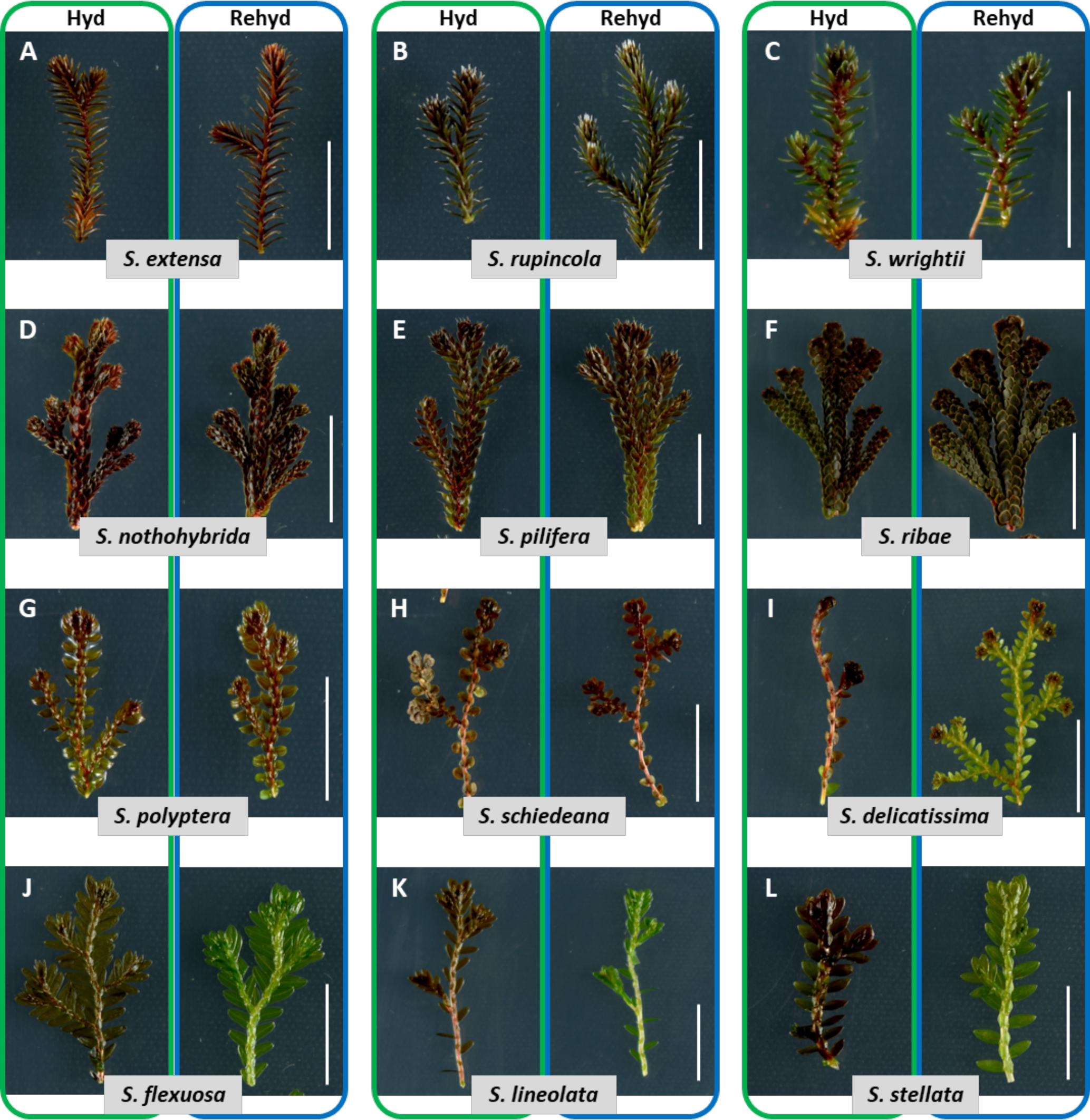
Viability test to determine desiccation tolerance in several *Selaginella* species. Tissue viability of explants exposed to desiccation and subsequent rehydration (Rehyd) was determined by the triphenyltetrazolium (TTC) test . Similar staining to tissue in hydrated (Hyd) conditions indicated that tissue is viable (red color). Desiccation-tolerant species (A – H), tissue-specific tolerant species (I), and desiccation-sensitive species (J – K). Scale bar = 1 cm

To corroborate that the explant dehydration system combined with TTC test can correctly identify desiccation-tolerant and sensitive species, we selected a tolerant and a sensitive species from those identified by the explant assay (*S. polyptera* and *S. flexuosa*), to be tested using the traditional whole plant desiccation assay. Plants were dehydrated by withholding water under greenhouse conditions for a period of 30 days (enough time to obtain a constant weight) and undergoing subsequent rehydration for recovery evaluation (**Fig. 6**). Individuals of *S. polyptera* showed the expected morphological changes in response to water loss (specifically stem curling), a common characteristic of desiccation-tolerant species (**Fig. 6B**). Tolerant individuals retained their green color during recovery evaluation, whereas the *S. flexuosa* individuals showed evident damage (tissue oxidation) and loss of turgidity during the recovery evaluation. Whole plant experiments were carried out for the remaining *Selaginella* samples and corroborated DT capacity of the following species: *S. extensa*, *S. nothohybrida*, *S. polyptera*, *S. ribae*, *S. rupincola*, *S. schiedeana*, and *S. wrightii* (**Supplementary Fig. S3**). However, the whole plant analysis may incorrectly classify a specimen such as *S. delicatissima* as a sensitive species because almost all tissue showed excessive oxidation (**Supplementary Fig. S3**). In contrast, the TTC assay had the advantage of indicating that this species possesses a DT mechanism specific to apical tissue (**Fig. 5I**).

**Figure 6.**
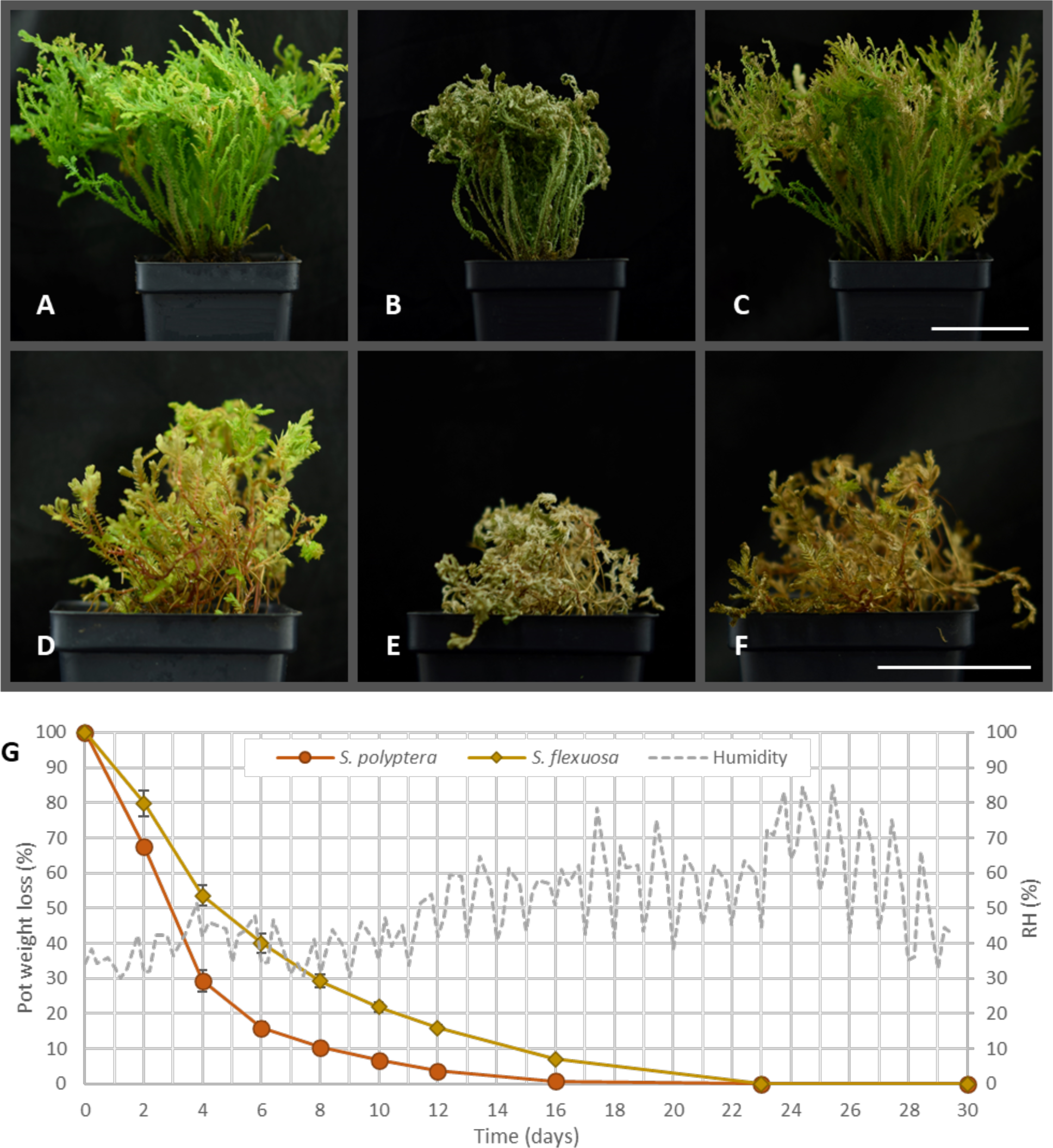
Evaluation of desiccation tolerance at the whole plant level. Whole plant analysis of *S. polyptera* (A-C) and *S. flexuosa* (D-F) under fully hydrated conditions (A, D), desiccated state (1 month without water) (B, E), and 48 h after rehydration (C, F). (G) Pot weight loss and greenhouse records of mean relative humidity (RH) during desiccation treatment. Each point represents the mean value of 3 replicates (pots), and error bars indicate standard deviation. Scale bar = 5 cm

### Loss of tissue viability at specific water contents

The viability of *S. silvestris* during dehydration was determined at different levels of water content using the TTC assay. Fully hydrated explants (full turgor) were dehydrated with the drying rate produced by MgCl_2_ and at specific relative water content (RWC) explants were removed from the drying system and immediately rehydrated. An initial analysis of tissue viability assayed by TTC staining revealed a change in viability between explants dehydrated to 40 and 20% RWC (**Fig. 7A**). The proportion of viable tissue was determined by image analysis of the red stained areas within explants (**Fig. 7B**). A detailed analysis at these water contents determined that the viable area of explants decreased by 13.5% at 40% RWC and further decreased by 67.8% at 20% RWC (**Fig. 7C**). Although most of the explant area remains viable at 40% RWC, measurement of quantum efficiency indicated that the tissue was under stress with a Fv/Fm=0.53 (**Supplementary Fig. S4**) compared to 0.8 in unstressed tissue in both desiccation-tolerant and sensitive species (**Fig. 2B**). In addition, quantum efficiency showed an even lower value at 20% RWC (Fv/Fm=0.47; **Supplementary Fig. S4**).

**Figure 7.**
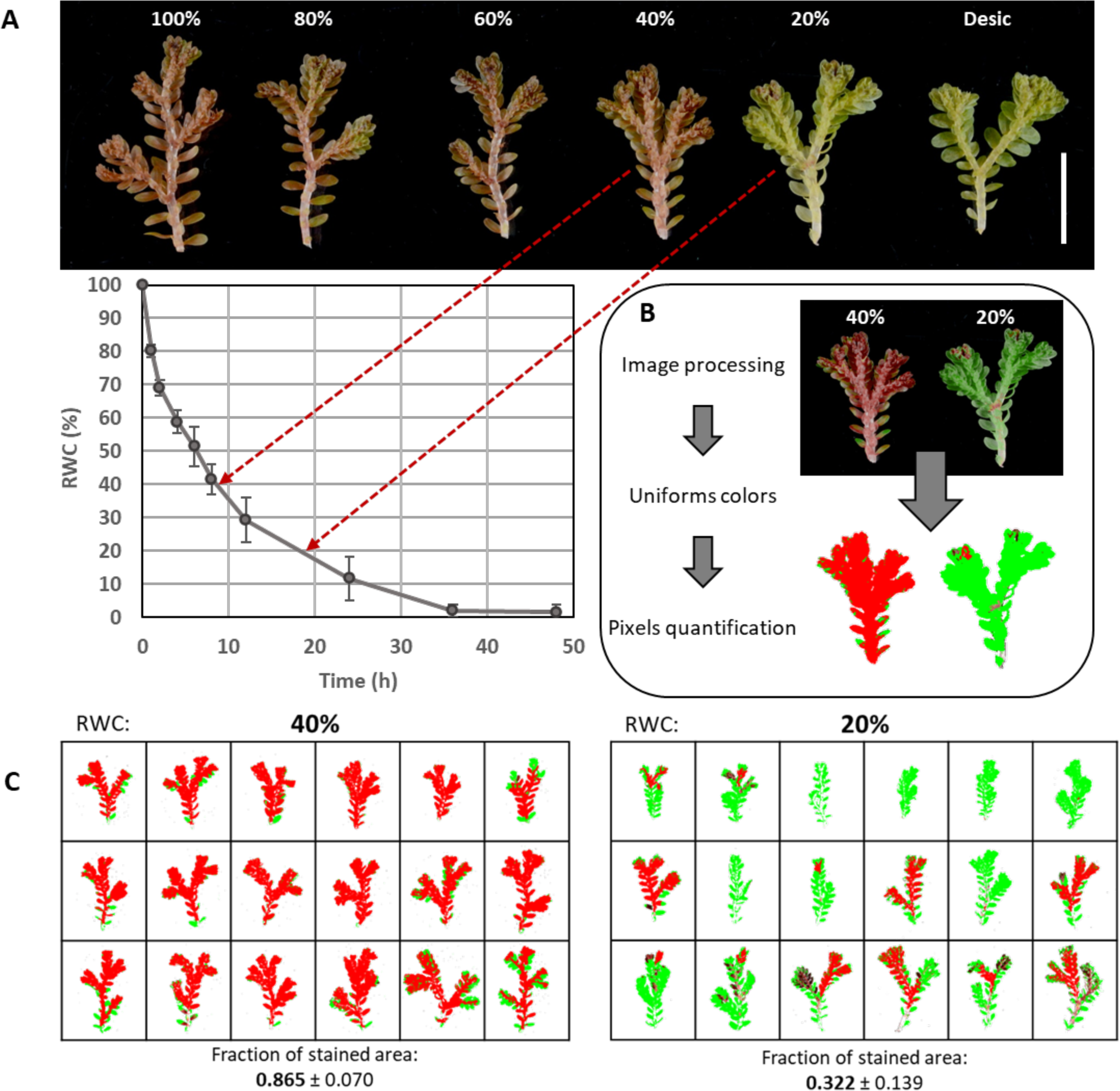
Loss of viability in *S. silvestris* explants. (A) Explant drying at specific relative water contents (RWC) and respiratory chain activity determination through triphenyltetrazolium test. (B) Main steps of the image analysis for determination of viable area (stained tissue) at each RWC. (C) Proportion of viable area of explants dehydrated to 40 and 20% RWC. Each of the points of the drying curve correspond to 6 different replicates, error bars represent standard deviation. Scale bar = 1 cm

Desiccation-sensitive *Selaginella* species such as *S. denticulata* and *S. silvestris* can be easily propagated via explants, the ratio of explants successfully established in substrate (that can create a new individual) to total explants is close to 1 in these species. Explants of *S. silvestris* dehydrated to 80 and 60% RWC can recover and re-establish growth, indicating that if tissue damage occurred at these water contents it was limited to a repairable level. At 40% RWC most of the explant area is viable according to the TTC test (**Fig. 7C**), but not all the explants were capable of continued growth in substrate and most of them had large sections that suffered visible damage (**Supplementary Fig. S4**). After 4 weeks in substrate, explants that survived dehydration to 40% RWC showed green sectors with clear growth and the emergence of at least one rhizophore (a specialized structure which produces roots). The failure of re-establish growth of some explants (around 18%) suggests that irreversible damage begins when the tissue is dehydrated to 40% RWC. Moreover, the TTC staining pattern of *S. silvestris* at 20% RWC revealed that in some explants the apical region was still viable, which also indicated that the apical region was the last part of the explant to lose viability. However, these small viable sectors showed a much slower growth (limited to a green tip and only few developed rhizophores) compared to explants dehydrated to 40% RWC (**Supplementary Fig. S4**). Our results indicated 40% RWC as the onset of a critical threshold during dehydration for *S. silvestris* that already causes irreversible damage in large sectors of most explants.

## Discussion

The dehydration techniques and methods of assessing recovery used for plant DT studies can be quite variable complicating cross-species comparisons. Desiccation experiments are commonly performed at the whole plant level under greenhouse or growth chamber conditions. However, whole plant procedures rely on carefully controlled conditions (especially humidity) that have a direct effect on the drying rate and thus the reproducibility of the experiments. Experiments using excised tissue inside a small, closed container with a dehydration agent constitute a simple and better controlled way to expose tissue of different species to the same drying conditions. Additional advantages of this drying technique include a reduced evaluation period and the potential for use under field conditions which would avoid the removal of the individual from its habitat. The use of excised tissues to perform comparative desiccation studies can be applied to homoiochlorophyllous species, where detached tissues retain DT (Dinakar *et al*., 2012; Mitra *et al*., 2013). Indeed, the homoiochlorophyllous strategy is estimated to comprise the majority of desiccation-tolerant plants (Tuba *et al*., 1998; Marks *et al*., 2021).

Although DT in *Selaginella* involves some constitutive protection components, species of this genus also require time to activate protection mechanisms induced during water loss (Liu *et al*., 2008; Yobi *et al*., 2012). Leakage measurements revealed that explants exposed to rapid drying showed significant membrane damage in *S. lepidophylla* and that moderate drying rates produced lower membrane damage in tolerant *Selaginella* species. Contrary to the assumption that a slow drying rate would produce the least damage, our results showed the slow regime can result in higher electrolyte leakage than moderate drying. This is probably the result of greater damage occurred during prolonged exposure to intermediate water contents during slower dehydration rates. One possible scenario stems from the observation that carbon fixation in desiccation-tolerant species ceases around -2 MPa, but light-harvesting reactions of photosynthesis continue until -15 MPa (Oliver *et al*., 2020) producing high-energy intermediates that can result in the production of toxic levels of ROS.

Measurement of the photosynthetic parameter Fv/Fm has previously been used to determine recovery after desiccation for *Selaginella* (Pandey *et al*., 2010; Xu *et al*., 2018; Alejo-Jacuinde *et al*., 2020). Upon rehydration, desiccation-tolerant species completely recovered Fv/Fm values after rehydration, whereas sensitive species exhibited values similar to the desiccated state indicating that the tissues were still under considerable stress. During rehydration chlorophyll contents also almost fully recovered in desiccation-tolerant species but not in sensitive species. Our results are in agreement with those previously reported showing that the desiccation-tolerant species *S. bryopteris* showed a similar pattern of Fv/Fm and chlorophyll content in response to desiccation (Pandey *et al*., 2010). The tolerant species, *S. lepidophylla* and *S. sellowii*, can tolerate and recover from the leakage of 22.7 and 14.1% of its total electrolytes, respectively but membrane integrity is completely compromised upon rehydration for the desiccation-sensitive species *S. silvestris* and *S. denticulata* (REL around 90%). These results with explants are comparable to those obtained for whole plants of *Selaginella* species (Agduma and Sese, 2016).

Maintaining the integrity of cellular components such as ribosomes is essential to survive desiccation. A comparative analysis between species of the Linderniaceae family showed that levels of total and polysomal RNA from desiccation-tolerant plants are maintained during the whole dehydration treatment, whereas a sensitive species showed RNA degradation at late dehydration times (Juszczak and Bartels, 2017). In our experiments, the integrity of rRNA was maintained in desiccation-tolerant *Selaginella* species during desiccation and rehydration processes, while in sensitive species rRNA integrity was maintained in the desiccated state but significant rRNA degradation was detected during rehydration. Cells undergo mechanical, structural and metabolic stresses during water loss (Oliver *et al*., 2020), but cellular damage can also occur during the influx of water at rehydration (Alpert and Oliver, 2002; Oliver *et al*., 2005). The precise stage at which most of the cellular damage occurs (during dehydration or rehydration) and whether cellular components differ in their tolerance capacity still remain uncertain.

Excessive ROS accumulation results in cell death, therefore, antioxidant protection systems play an essential role in DT (Proctor and Tuba, 2002; Challabathula *et al*., 2018; Oliver *et al*., 2020). In fact, maintenance of antioxidant potential in desiccation-tolerant organisms, particularly glutathione redox potential, has been proposed as a marker of viability during long-term desiccation (Kranner and Birtić, 2005). Antioxidant potential in *Selaginella* explants (determined by FRAP and DPPH assays), was greater in desiccation-tolerant than sensitive species under hydrated conditions. High antioxidant levels under normal conditions suggest that *S. lepidophylla* and *S. sellowii* are primed to tolerate forthcoming desiccation events, as described for the DT grass *Sporobolus stapfianus* (Oliver *et al*., 2011). Antioxidant capacity increased in all *Selaginella* species in response to desiccation, although desiccation-tolerant plants accumulate antioxidants to a greater extent. Antioxidants are also required to scavenge ROS at the rehydration stage (Farrant *et al*., 2007; Oliver *et al*., 2020) which is consistent with our observation that desiccation-tolerant *Selaginella* species maintained higher antioxidant potential during rehydration. Other antioxidant compounds like polyphenols and flavonoids have a putative membrane protective role during desiccation (Georgieva *et al*., 2017). Metabolic analysis of hydrated and dehydrated plants determined a higher level of several amino acid-derived flavonoids in *S. lepidophylla* compared with the sensitive species *S. moellendorffii* (Yobi *et al*., 2012). Additionally, a higher fraction of genes involved in flavonoid metabolism is differentially expressed in response to dehydration in *S. lepidophylla* and *S. sellowii* compared to *S. denticulata* (Alejo-Jacuinde *et al*., 2020). This is consistent with our findings that tolerant plants showed higher levels of flavonoids in hydrated conditions and accumulated more during desiccation than sensitive species.

Our results also provide valuable information for tolerant species comparisons. Although both *S. lepidophylla* and *S. sellowii* have high antioxidant levels at the desiccated stage, in response to desiccation *S. lepidophylla* showed a significant increase in antioxidant potential, whereas *S. sellowii* explants showed only a slight increase in the antioxidant parameters. These results suggest that antioxidant protection in *S. lepidophylla* is inducible whereas in *S. sellowii* is largely constitutive. The ability to survive rapid desiccation that exhibit some bryophytes is associated to a constitutive protection mechanism (Oliver *et al*., 2000; Alpert and Oliver, 2002), which also showed a rapid recovery of photosynthesis (Challabathula *et al*., 2018). A major constitutive component in the DT in *S. sellowii* correlates with a lower membrane damage induced by different drying rates and faster recovery of photosynthesis during rehydration compared to *S. lepidophylla* (Alejo-Jacuinde *et al*., 2020).

The TTC test was successfully adapted to vegetative tissue of *Selaginella* providing a simple identification of viable tissue after desiccation treatment. The efficiency of the TTC assay as viability marker for DT was shown with the identification of novel desiccation-tolerant species. We were able to establish that *S. extensa*, *S. nothohybrida*, *S. polyptera*, *S. ribae*, *S. rupincola*, *S. schiedeana*, and *S. wrightii* are desiccation-tolerant species. Previous studies had described *S. nothohybrida* and *S. rupincola* as “resurrection species” (Aguilar *et al*., 2015; Yu *et al*., 2017) but no data delineating its DT capacity was reported. This assay also revealed that only the apices of the explants of *S. delicatissima* exhibited DT which could enable this species to re-establish growth and create a new individual after an extensive dry period. This sample was classified as a tissue-specific tolerant species, further studies on this *Selaginella* specimen could reveal insights into why the apical tissue has the capability for DT.

Interestingly, *S. extensa*, a desiccation-tolerant species with isophyllous morphology, was collected from a population located in a cloud forest with an annual rainfall of around 2028 mm. Most of the isophyllous *Selaginella* species occupy xeric habits (Arrigo *et al*., 2013) and at least eight of these have been classified as desiccation-tolerant (Proctor and Pence, 2002). We identified three additional desiccation-tolerant isophyllous species (*S. extensa*, *S. rupincola*, and *S. wrightii*). All these species belong to the same clade within the *Selaginella* phylogeny (Homoeophyllae according to Zhou *et al*., 2015; Rupestrae according to Weststrand and Korall, 2016), suggesting that all species within this clade are likely desiccation-tolerant (between 45 – 60 species), including those that naturally grow in very moist habitats.

A whole plant study reported that the sensitive species *S. moellendorffii* cannot recover from dehydration to 40% RWC or below (Yobi *et al*., 2012). Others have suggested that dehydration below 40% RWC produces extensive cellular damage resulting in death of most of the plants (Mitra *et al*., 2013). An extensive literature analysis by Zhang and Bartels (2018) proposed a boundary between dehydration and the desiccation response at 40 – 30% RWC. Our data confirmed that viability loss initiates at 40% RWC in the sensitive species *S. silvestris* already causing extensive irreversible damage in large portion of the explants and the majority of the explant did not survive below 40% RWC.

The use of explants in desiccation experiments has several advantages, and our results corroborate that the response to desiccation in explants is similar to that observed in whole plant analysis. Our main findings are summarized in **Figure 8**, which also indicated the methodologies proposed to evaluate each characteristic associated to DT. The system discussed here can be exploited for the characterization of desiccation-tolerant species and to compare tolerance mechanisms between them. The main viability marker proposed in this study (TTC), represents a simple and robust method to determine DT capacity. Our data also contributes to the description of perhaps a distinct tissue-specific DT mechanism in *Selaginella* in the apical portion of the tissue.

**Figure 8.**
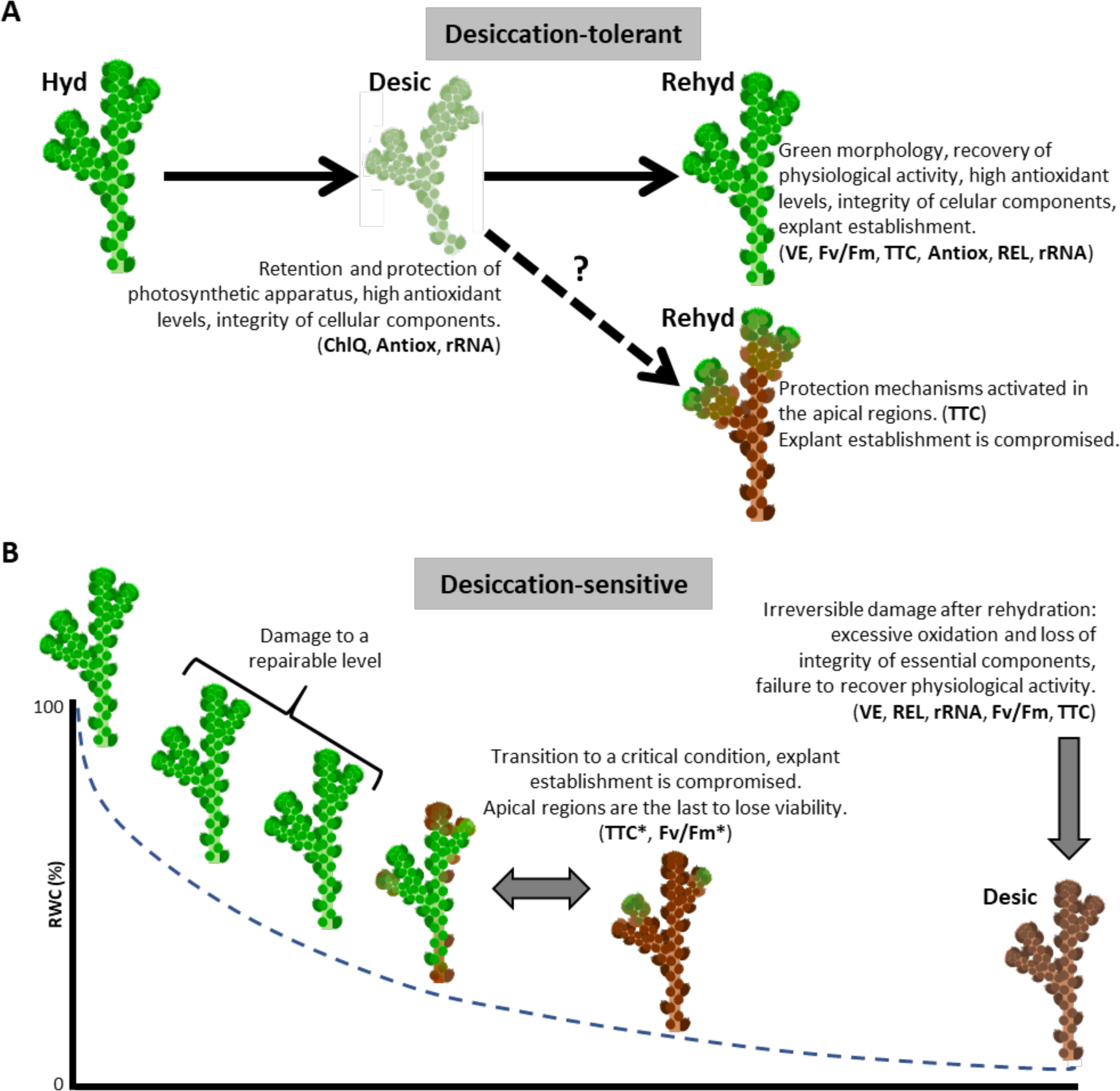
Desiccation tolerance associated characteristics and critical stage determination in *Selaginella* species. (A) Overview of measurable responses, components, or activities in desiccation-tolerant species. Tissue-specific desiccation tolerance indicated as an alternative mechanism (dashed arrow). (B) Identification of a critical condition by viability loss in sensitive species. Methodologies proposed to evaluate each characteristic are indicated in bold (in brackets): antioxidant potential (Antiox), chlorophyll quantification (ChlQ), maximum quantum efficiency of PSII (Fv/Fm), relative electrolyte leakage (REL), rRNA integrity (rRNA), triphenyltetrazolium chloride (TTC), visual evaluation (VE). Tissue dehydrated and subsequently rehydrated (*).

## Supplementary data

Supplementary data are available at *JXB* online.

*Supplementary Table S1*. Collection sites of *Selaginella* species included in the study.

*Supplementary Protocol S1*. Drying system, biological material, TTC test and imagen analysis of viable areas.

*Supplementary Fig. S1*. Drying curve of *Selaginella* species.

*Supplementary Fig. S2*. Chlorophyll retention after desiccation process in tolerant species.

*Supplementary Fig. S3*. Whole plant level evaluation of desiccation tolerance in several *Selaginella* species.

*Supplementary Fig. S4*. Recovery of *S. silvestris* explants at specific water contents.

## Acknowledgements

We are thankful to Araceli Oropeza-Aburto and Katia Gil-Vega for their technical assistance. We also thank Dr. Andrés Estrada-Luna for his technical support and lending us the chlorophyll fluorimeter. We are grateful to Dalia Grego-Valencia for her help in the morphological identification of specimens. G.A.-J. is indebted to CONACyT (Mexico) for PhD fellowship. This work was supported in part by grants from the Basic Science program from CONACytT (Grant 00126261), the Governor University Research Initiative program (05-2018) from the State of Texas, and by a Senior Scholar grant from Howard Hughes Medical Institute (grant 55005946) to L.H.-E.

## Author contributions

G.A.-J. and L.H.-E. designed the research. G.A.-J. conducted most of the experimental research. G.A.-J. and T.K.-G. performed electrolyte leakage analysis. N.M.-G. and J.P.D.-F. carried out antioxidant analysis. G.A.-J., T.K.-G., K.M. and D.T.-D. obtained plant material and classified novel tolerant species. G.A.-J., L.H.-E., J.S. and M.O. analyzed the data. G.A.-J., J.S. and L.H.-E. wrote the paper. All authors read and approved the manuscript.

## Data availability

All data supporting the findings of this study are available within the paper and within its supplementary materials published online.

## References

Agduma AR, Sese MD. 2016. Cellular Biochemical Changes in Selaginella tamariscina (Beauv.) Spring and Sellaginella plana (Desv. ex Poir.) Heiron. as Induced by Desiccation. Tropical life sciences research 27, 37–52.

Aguilar MI, Benítez W V, Colín A, Bye R, Ríos-Gómez R, Calzada F. 2015. Evaluation of the diuretic activity in two Mexican medicinal species: Selaginella nothohybrida and Selaginella lepidophylla and its effects with ciclooxigenases inhibitors. Journal of ethnopharmacology 163, 167–72.

Alejo-Jacuinde G, González-Morales SI, Oropeza-Aburto A, Simpson J, Herrera-Estrella L. 2020. Comparative transcriptome analysis suggests convergent evolution of desiccation tolerance in Selaginella species. BMC Plant Biology 20, 1–18.

Alpert P. 2005. The limits and frontiers of desiccation-tolerant life. Integrative and comparative biology 45, 685–95.

Alpert P, Oliver MJ. 2002. Drying without dying. In: Black M,, In: Pritchard HW, eds. Desiccation and survival in plants: drying without dying. Wallingford: CABI, 3–43.

Arrigo N, Therrien J, Anderson CL, Windham MD, Haufler CH, Barker MS. 2013. A total evidence approach to understanding phylogenetic relationships and ecological diversity in Selaginella subg. Tetragonostachys. American Journal of Botany 100, 1672–1682.

Bartels D, Hussain SS. 2011. Resurrection Plants: Physiology and Molecular Biology. Springer Berlin Heidelberg, 339–364.

Bewley JD. 1979. Physiological aspects of desiccation tolerance. Annual Review of Plant Physiology 30, 195–238.

Challabathula D, Puthur JT, Bartels D. 2016. Surviving metabolic arrest: photosynthesis during desiccation and rehydration in resurrection plants. Annals of the New York Academy of Sciences 1365, 89–99.

Challabathula D, Zhang Q, Bartels D. 2018. Protection of photosynthesis in desiccation-tolerant resurrection plants. Journal of Plant Physiology 227, 84–92.

Dinakar C, Bartels D. 2013. Desiccation tolerance in resurrection plants: new insights from transcriptome, proteome and metabolome analysis. Frontiers in plant science 4, 482.

Dinakar C, Djilianov D, Bartels D. 2012. Photosynthesis in desiccation tolerant plants: energy metabolism and antioxidative stress defense. Plant science : an international journal of experimental plant biology 182, 29–41.

Easlon HM, Bloom AJ. 2014. Easy Leaf Area: Automated Digital Image Analysis for Rapid and Accurate Measurement of Leaf Area. Applications in Plant Sciences 2, 1400033.

Eickmeier WG. 1980. Photosynthetic Recovery of Resurrection Spikemosses from Different Hydration Regimes.

Farrant JM, Brandt W, Lindsey GG. 2007. An Overview of Mechanisms of Desiccation Tolerance in Selected Angiosperm Resurrection Plants. Plant Stress 1, 72–84.

Georgieva K, Dagnon S, Gesheva E, Bojilov D, Mihailova G, Doncheva S. 2017. Antioxidant defense during desiccation of the resurrection plant Haberlea rhodopensis. Plant Physiology and Biochemistry 114, 51–59.

Hoekstra FA, Golovina EA, Buitink J. 2001. Mechanisms of plant desiccation tolerance. Trends in Plant Science 6, 431–438.

Juszczak I, Bartels D. 2017. LEA gene expression, RNA stability and pigment accumulation in three closely related Linderniaceae species differing in desiccation tolerance. Plant Science 255.

Kranner I, Birtić S. 2005. A modulating role for antioxidants in desiccation tolerance. Integrative and Comparative Biology 45, 734–740.

Lebkuecher JG, Eickmeier WG. 1991. Reduced photoinhibition with stem curling in the resurrection plant Selaginella lepidophylla. Oecologia 88, 597–604.

Leprince O, Buitink J. 2015. Introduction to desiccation biology: from old borders to new frontiers. Planta 242, 369–378.

Lin CH, Chen BS, Yu CW, Chiang SW. 2001. A water-based triphenyltetrazolium chloride method for the evaluation of green plant tissue viability. Phytochemical Analysis 12, 211–213.

Liu M-S, Chien C-T, Lin T-P. 2008. Constitutive components and induced gene expression are involved in the desiccation tolerance of Selaginella tamariscina. Plant & cell physiology 49, 653–63.

Lopez Del Egido L, Navarro-Miró D, Martinez-Heredia V, Toorop PE, Iannetta PPM. 2017. A spectrophotometric assay for robust viability testing of seed batches using 2,3,5-triphenyl tetrazolium chloride: Using Hordeum vulgare L. as a model. Frontiers in Plant Science 8, 747.

Maranz S, Wiesman Z, Garti N. 2003. Phenolic constituents of shea (Vitellaria paradoxa) kernels. Journal of Agricultural and Food Chemistry 51, 6268–6273.

Marks RA, Farrant JM, Nicholas McLetchie D, VanBuren R. 2021. Unexplored dimensions of variability in vegetative desiccation tolerance. American Journal of Botany, ajb2.1588.

Mickel JT, Smith AR. 2004. The Pteridophytes of Mexico. Part I (Descriptions and Maps). New York: The New York Botanical Garden Press.

Mitra J, Xu G, Wang B, Li M, Deng X. 2013. Understanding desiccation tolerance using the resurrection plant Boea hygrometrica as a model system. Frontiers in Plant Science 4, 446.

Oliver MJ, Farrant JM, Hilhorst HWM, Mundree S, Williams B, Bewley JD. 2020. Desiccation Tolerance: Avoiding Cellular Damage During Drying and Rehydration. Annual Review of Plant Biology 71.

Oliver MJ, Guo L, Alexander DC, Ryals JA, Wone BWM, Cushman JC. 2011. A Sister Group Contrast Using Untargeted Global Metabolomic Analysis Delineates the Biochemical Regulation Underlying Desiccation Tolerance in *Sporobolus stapfianus*. The Plant Cell 23, 1231–1248.

Oliver MJ, Tuba Z, Mishler BD. 2000. The evolution of vegetative desiccation tolerance in land plants. Plant Ecology 151, 85–100.

Oliver MJ, Velten J, Mishler BD. 2005. Desiccation Tolerance in Bryophytes: A Reflection of the Primitive Strategy for Plant Survival in Dehydrating Habitats? Integrative and Comparative Biology 45, 788–799.

Pandey V, Ranjan S, Deeba F, Pandey AK, Singh R, Shirke PA, Pathre U V. 2010. Desiccation-induced physiological and biochemical changes in resurrection plant, Selaginella bryopteris. Journal of Plant Physiology 167, 1351–1359.

Proctor MCF, Pence VC. 2002. Vegetative tissues: bryophytes, vascular resurrection plants and vegetative propagules. In: Black M,, In: Pritchard HW, eds. Desiccation and survival in plants: drying without dying. Wallingford: CABI, 207–237.

Proctor MCF, Tuba Z. 2002. Poikilohydry and homoihydry: antithesis or spectrum of possibilities? New Phytologist 156, 327–349.

Richardson AD, Duigan SP, Berlyn GP. 2002. An evaluation of noninvasive methods to estimate foliar chlorophyll content. New Phytologist 153, 185–194.

Ruf M, Brunner I. 2003. Vitality of tree fine roots: reevaluation of the tetrazolium test. Tree physiology 23, 257–63.

Sakanaka S, Tachibana Y, Okada Y. 2005. Preparation and antioxidant properties of extracts of Japanese persimmon leaf tea (kakinoha-cha). Food Chemistry 89, 569–575.

Tuba Z, Protor MCF, Csintalan Z. 1998. Ecophysiological responses of homoiochlorophyllous and poikilochlorophyllous desiccation tolerant plants: a comparison and an ecological perspective.

Weststrand S, Korall P. 2016. Phylogeny of Selaginellaceae: There is value in morphology after all! American journal of botany 103, 2136–2159.

Xiao L, Yobi A, Koster KL, He Y, Oliver MJ. 2018. Desiccation tolerance in *Physcomitrella patens* : Rate of dehydration and the involvement of endogenous abscisic acid (ABA). Plant, Cell & Environment 41, 275–284.

Xu Z, Xin T, Bartels D, et al. 2018. Genome Analysis of the Ancient Tracheophyte Selaginella tamariscina Reveals Evolutionary Features Relevant to the Acquisition of Desiccation Tolerance. Molecular Plant 11, 983–994.

Yahia EM, Gutierrez-Orozco F, Leon CA de. 2011. Phytochemical and antioxidant characterization of the fruit of black sapote (Diospyros digyna Jacq.). Food Research International 44, 2210–2216.

Yobi A, Wone BWM, Xu W, Alexander DC, Guo L, Ryals JA, Oliver MJ, Cushman JC. 2012. Comparative metabolic profiling between desiccation-sensitive and desiccation-tolerant species of Selaginella reveals insights into the resurrection trait. Plant Journal 72, 983–999.

Yu R, Baniaga AE, Jorgensen SA, Barker S, American S, Journal F. 2017. A Successful in vitro Propagation Technique for Resurrection Plants of the Selaginellaceae A Successful in vitro Propagation Technique for Resurrection Plants of the Selaginellaceae. American Fern Journal 107, 96–104.

Zhang Q, Bartels D. 2018. Molecular responses to dehydration and desiccation in desiccation-tolerant angiosperm plants. Journal of Experimental Botany 69, 3211–3222.

Zhang Q, Song X, Bartels D. 2016. Enzymes and Metabolites in Carbohydrate Metabolism of Desiccation Tolerant Plants. Proteomes 4.

Zhou XM, Rothfels CJ, Zhang L, et al. 2015. A large-scale phylogeny of the lycophyte genus Selaginella (Selaginellaceae: Lycopodiopsida) based on plastid and nuclear loci. Cladistics 32, 360–389.

